# Extraction and quantification of lineage-tracing barcodes with NextClone and CloneDetective

**DOI:** 10.1101/2023.11.19.567755

**Authors:** Givanna H. Putri, Nichelle Pires, Nadia M. Davidson, Catherine Blyth, Aziz M. Al’Khafaji, Shom Goel, Belinda Phipson

**Affiliations:** The Walter and Eliza Hall Institute of Medical Research and The Department of Medical Biology, The University of Melbourne, Parkville, VIC, Australia; Peter MacCallum Cancer Centre and The Sir Peter MacCallum Department of Oncology, University of Melbourne, Parkville, VIC, Australia; Broad Institute of MIT and Harvard, Cambridge, Massachusetts, USA

## Abstract

**Summary:** The study of clonal dynamics has significantly advanced our understanding of cellular heterogeneity and lineage trajectories. With recent developments in lineage-tracing protocols such as ClonMapper or SPLINTR, which combine DNA barcoding with single-cell RNA sequencing (scRNA-seq), biologists can trace the lineage and evolutionary paths of individual clones while simultaneously observing their transcriptomic changes over time. Here, we present NextClone and CloneDetective, an integrated highly scalable Nextflow pipeline and R package for efficient extraction and quantification of clonal barcodes from scRNA-seq data and DNA sequencing data tagged with lineage-tracing barcodes. We applied both NextClone and CloneDetective to data from a barcoded MCF7 cell line and demonstrate their utility for advancing clonal analysis in the era of high-throughput sequencing.

**Availability and implementation:** NextClone and CloneDetective are freely available and open-source on github (https://github.com/phipsonlab/NextClone and https://github.com/phipsonlab/CloneDetective). Documentations and tutorials for NextClone and CloneDetective can be found at https://phipsonlab.github.io/NextClone/ and https://phipsonlab.github.io/CloneDetective/respectively.

## Introduction

Recent advances in lineage-tracing protocols, such as ClonMapper [1], SPLINTR [2], Larry [3] and LoxCode [4], which integrate DNA barcoding with single-cell RNA sequencing (scRNA-seq), have catalysed significant progress in clonal biology. These techniques allow biologists to explicitly trace the ancestry and evolutionary trajectories of individual clones while observing their transcriptional changes over time.

Despite these methodological advances, there is a notable gap in the development of computational tools to efficiently extract and quantify clonal barcodes from scRNA-seq data and DNA barcoding data (DNA-seq data) which exclusively sequences clone barcode reads using Next Generation Sequencing.

To address this, we present NextClone and CloneDetective: a Nextflow [5] pipeline and R package designed specifically for extracting and quantifying clonal barcodes from both barcoded DNA-seq and scRNA-seq data. We demonstrate the utility and efficacy of NextClone and CloneDetective using DNA-seq and scRNA-seq data from the MCF7 breast cancer cell line, barcoded using the ClonMapper lineage tracing protocol [1].

## Design and Usage

### Core principle of NextClone

NextClone is a pipeline developed using Nextflow, an open-source workflow manager that simplifies the creation and deployment of scalable reproducible analysis workflows. It is designed with scalability in mind, to facilitate rapid extraction and quantification of clonal barcodes from both DNA-seq and scRNA-seq data.

The pipeline comprises two distinct workflows (Figure 1A), one for DNA-seq data and the other for scRNA-seq data. Both workflows are highly modular and adaptable, with software that can easily be substituted as required, and with parameters that can be tailored through the *nextflow*.*config* file to suit diverse needs. The architecture of NextClone takes full advantage of Nextflow’s task execution caching and checkpointing mechanisms, allowing for software and parameter adjustments without the need to rerun the unaffected parts of the workflow.

**Figure 1.**
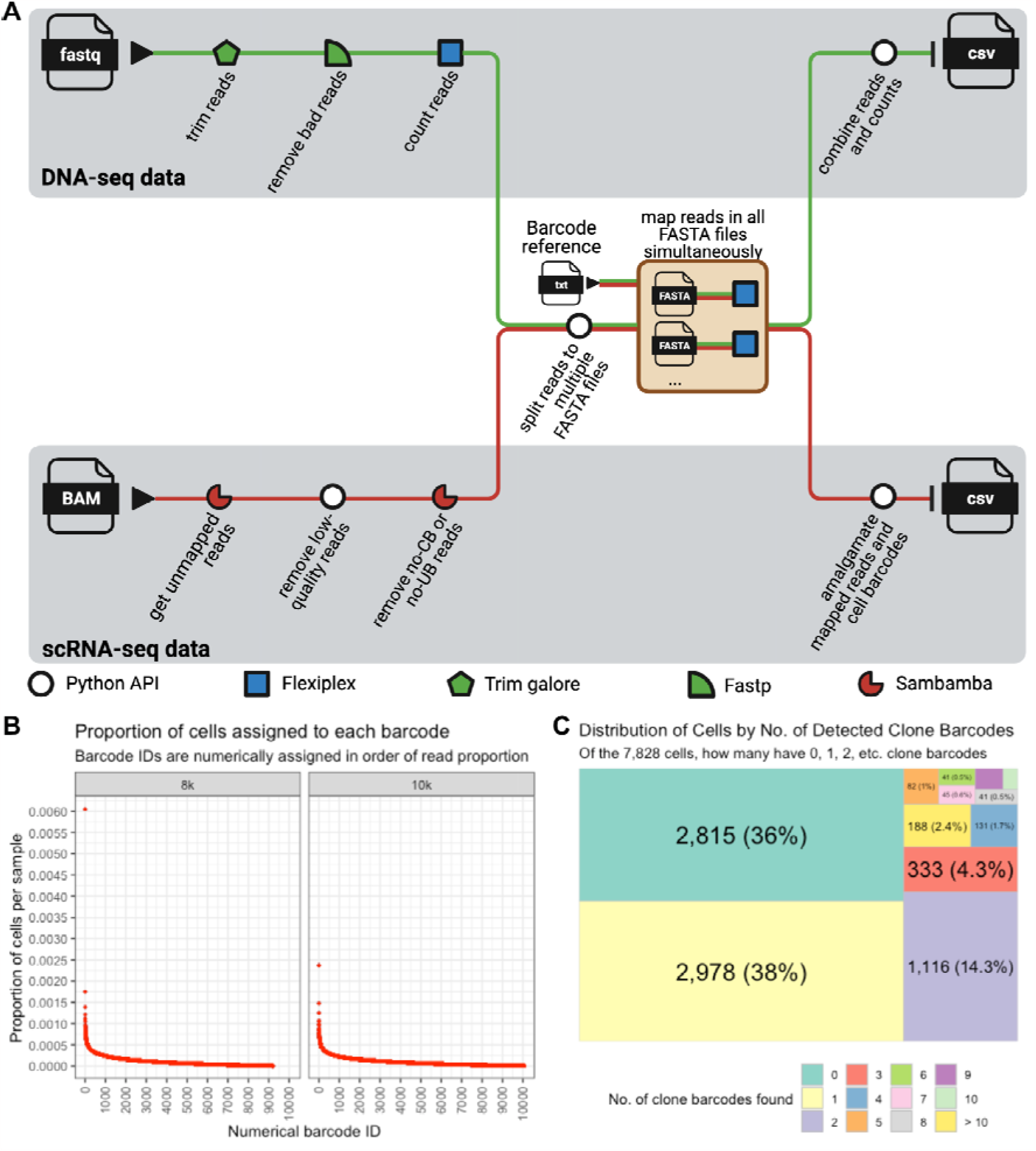
Overview of NextClone pipeline and example visualisations from the CloneDetective R Package. (A) Overview of NextClone pipeline for extracting and quantifying clonal barcodes in DNA-seq data and scRNA-seq data. (B) An ordered clonal abundance plot illustrating the distribution of the clone abundance (defined as the proportion of cells in each clone) for 8k and 10k DNA-seq data. The proportion is computed against the total number of cells in the sample. (C) A treemap visualising the distribution of cells (as absolute number and proportion) according to the number of detected clone barcodes within each cell for the scRNA-seq data from the barcoded MCF7 cell line. The treemap categorises cells into groups based on the number of clone barcodes detected per cell, ranging from cells with no detected barcodes to cells with ten or more barcodes detected. The grouping of cells was performed after removing reads resulting from PCR amplification.

Optimised for high-performance computing (HPC) platforms, NextClone comes with predefined process selectors (implemented using Nextflow’s *withLabel* selectors) which allocate computing resources effectively to each step, enhancing its scalability. These selectors, along with the computing resources, can be adjusted via a configuration file to fit various HPC configurations and computational needs.

Integral to the operation of NextClone are two key components: Flexiplex [6], a lightweight sequence search and demultiplexing tool, and a bespoke Python package which supports the parallel processing of samples. The latter capitalises on the vast computational resources of HPC platforms to expedite clone barcode extraction and quantification processes.

### NextClone DNA-seq workflow

Figure 1A illustrates the NextClone workflow for processing DNA-seq data. The workflow begins with trimming reads to a uniform length (by default 20 base pairs) using Trim Galore [7], [8], and removing low quality reads using Fastp [9]. Reads with over 20% of their nucleotides with Phred quality scores below 30 are removed. Users have the flexibility to modify these default settings to suit their experimental needs. Trim Galore and Fastp are independent tools and must be installed separately to the workflow.

Following the initial processing, Flexiplex is then used to count the occurrences of each unique read sequence. In its default setting, NextClone configures Flexiplex to run on multiple threads (default 4) to speed up the counting process. The Python package then divides the reads into several FASTA files, which can be increased from the default two as needed. Each FASTA file then undergoes a mapping process against a reference list of clonal barcode sequences provided by the user using Flexiplex running on multiple threads (default 4). Each mapping task is submitted as an individual job to the HPC scheduler, allowing all the tasks to be processed simultaneously as resources permit.

Upon completion of these jobs, NextClone aggregates the mapping results with the initial read counts to determine the frequency of each clonal barcode in the samples. The initial counting of reads, prior to mapping, helps to reduce the total number of reads that need to be mapped by Flexiplex, thereby improving efficiency. The final output is a CSV file, where each row corresponds to a unique clone barcode and its frequency in the sample.

We applied NextClone and Pycashier [1], [10], an alternative pipeline which takes a clustering based approach, to extract clone barcodes from two DNA-seq datasets, namely the 8k and 10k datasets. These datasets were derived from an MCF7 breast cancer cell line barcoded with the ClonMapper library [1] and expanded into 8,000 and 10,000 cell libraries (see Supplementary Material for details of the datasets). Both pipelines detected similar numbers of barcodes for both DNA-seq datasets under different filtering thresholds, with NextClone detecting slightly fewer barcodes compared to Pycashier (Supplementary Material Figure S1A). NextClone presents a complementary method to Pycashier by mapping reads to a known list of barcodes, an approach that can enhance precision in identifying clonal barcodes. This direct mapping technique, used alongside or in contrast to clustering based approaches like Pycashier, can be beneficial in studies where precise barcode detection is crucial.

### NextClone scRNA-seq workflow

Figure 1A depicts the NextClone workflow for extracting and identifying the clone barcodes of cells in a barcoded scRNA-seq dataset. The workflow takes as input, an indexed and position-sorted Binary Alignment Map (BAM) file, typically generated by software such as 10x CellRanger [11] which aligns sequencing reads to a reference genome and performs feature counting to generate a cell-by-gene matrix from FASTQ files. The BAM file first undergoes a filtering process using Sambamba [12] to retain only unmapped reads, which are reads that could not be mapped to the genome. The Python package then removes lowquality reads, specifically those with a mean Phred score lower than a predefined threshold.

Reads missing the 10x cell barcodes or Unique Molecular Identifier (UMI) barcodes, denoted by the absence of CB and UB tags, are also discarded.

Similar to the DNA-seq workflow, the remaining unmapped reads are then divided into multiple FASTA files. To maximise efficiency, each file is mapped independently to a reference list of clone barcode sequences using Flexiplex, with each mapping task executed as a separate job on the HPC platform. The workflow culminates by amalgamating the mapped barcodes into a single CSV file where each row corresponds to a read associated with a 10x cell barcode, a UMI, and a clone barcode. This consolidated output facilitates subsequent analyses and enables the extraction of critical metrics such as clone barcode frequency and the number of distinct clone barcodes detected per cell, which are essential for downstream analysis.

### Using CloneDetective to Interrogate NextClone output for DNA-seq data

The CloneDetective R package contains functions to analyse clone barcode distributions from DNA-seq data which can inform the design of future scRNA-seq experiments. For a given set of experimental parameters (e.g. the number of cells captured), it can predict clone barcode representation in the resulting scRNA-seq data by estimating clonal abundance from DNA-seq data. To demonstrate its usage, we used CloneDetective to analyse the output of NextClone for the 8k and 10k DNA-seq datasets.

CloneDetective offers functions to count the number of clone barcodes in the dataset and to filter out clones with low abundance based on a predefined cell count threshold (Supplementary Material Figure S1A). It can also calculate the proportion of cells in each clone against the total number of cells in the sample and illustrate this as an ordered abundance plot (Figure 1B). Such plots are insightful for determining whether clones proliferated uniformly or if some clones were more dominant, potentially outcompeting and restricting the expansion of others.

Furthermore, CloneDetective uses these calculated cell proportions to forecast clonal abundance for future scRNA-seq experiments. This feature is helpful for estimating the number of cells to sequence for a given clonal library in order to maximise the chance of obtaining a reasonable representation of clones. For instance, using the clonal proportions derived from the 8k and 10k DNA-seq datasets (Figure 1B), it can estimate the distribution of clonal abundance when sequencing either 10,000 or 20,000 cells (Supplementary Material Figure S1B and S1C). These predictions are derived by scaling the clonal proportions to the total number of cells planned for sequencing. Additionally, CloneDetective can accommodate scenarios where clone barcodes are not detected in every single cell by incorporating binomial sampling into its estimations. CloneDetective can also estimate the number of clones that contain at least a specified minimum number of cells (Supplementary Material Figure S1D), as well as the cumulative number of cells for the top 200 most abundant clones (Supplementary Material Figure S1E).

Altogether, CloneDetective provides valuable predictive analytics for scRNA-seq experiment planning, ensuring comprehensive clone representation for meaningful downstream analysis.

### Using CloneDetective for barcoded scRNA-seq data

We developed functions within CloneDetective to perform an initial assessment of clone barcode uptake by cells, and to assign clone barcodes to cells. The resulting clone assignment can be exported to a CSV file, integrated into a cell-by-gene count matrix stored as SingleCellExperiment [13], AnnData [14], or Seurat object [15], and finally analysed using established scRNA-seq tools such as Scanpy [16], Seurat [15], and the suite of R packages in Bioconductor.

Ideally, a single clone barcode should be detected in each cell in an scRNA-seq experiment. However, it’s common to find cells with either no or multiple clone barcodes detected. In our analysis of the scRNA-seq data from the MCF7 cell lines barcoded using Clonmapper library [1] (see Supplementary Material for details on the dataset), we used CloneDetective to quantify the detection rate of clone barcodes and the proportion of cells containing specific numbers of clone barcodes—one, two, three, or more (Figure 1C). This information is essential for understanding the clone barcode uptake in the experiment.

To assign clone barcodes to cells, the first step is to consolidate duplicate reads from PCR amplifications. Reads are grouped based on their respective 10x cell barcodes and UMIs. If all reads in a group map to a single clone barcode, they are merged together. Conversely, if the reads map to multiple clone barcodes, they are combined and assigned to the clone barcode that represents at least 70% of the total reads in the group. This threshold can be adjusted to suit specific needs. Any groups that do not have a clone barcode meeting this criteria are discarded.

Upon UMI deduplication, we perform clone barcode assignment using the following stratified protocol. Cells with exactly one clone barcode are assigned to their respective clones. In our dataset, this scenario represents 38% of the cells in the sample (Figure 1C).

For cells with multiple clone barcodes detected, we resolved the assignment by evaluating the read abundance. Cells were first assigned to the clone barcode that accounted for over half of the detected PCR deduplicated reads. In our dataset, this approach effectively applied to 67% of such cells (and made up 26% of all the cells in the sample).

The remaining cells which did not meet the above criterion underwent further analysis based on average barcode edit distances. Here, cells were assigned to the clone barcode with the smallest average edit distance. In cases where distances were identical, the tie was broken in favour of the clone barcode with the higher read proportion.

## Discussion

NextClone and CloneDetective provide a comprehensive pipeline for extracting and quantifying clonal barcodes from DNA-seq and scRNA-seq data barcoded with lineage tracing protocols. They provide researchers with an effective framework for processing and interpreting DNA-seq and barcoded scRNA-seq data. NextClone is particularly engineered for high scalability to take full advantage of the vast computational resources offered by HPC platforms. CloneDetective is especially valuable for allowing researchers to use affordable DNA-seq experiments to guide the design of subsequent, more expensive scRNA-seq experiments. Both NextClone and CloneDetective are open-source and accessible on Github. We demonstrated the utility of NextClone and CloneDetective by analysing two DNA-seq and one scRNA-seq dataset of the MCF7 breast cancer cell line transduced with the ClonMapper library.

In the current workflows for both DNA-seq and scRNA-seq, NextClone identifies clone barcodes by mapping reads against a predefined list of barcodes. This allows NextClone to be adapted to work with any lineage tracing methodology, provided that a reference barcode list is available. Future work could implement a strategy for extracting and quantifying clonal barcodes when no known barcode list is present. Theoretically, Flexiplex could be used to construct such a list by cataloguing the most frequent read occurrences, which would then be used as the list of barcodes during the read mapping phase. This approach has been previously proven effective in long read single-cell sequencing [6], suggesting its high plausibility for use in this context where the barcodes are longer and have lower error rates. This approach could broaden the applicability of NextClone.

For scRNA-seq data, CloneDetective used a stratified protocol to assign clone barcodes to cells. Further work could investigate whether cells with equal read proportions for their two most abundant barcodes may represent doublets and whether clones can be identified that have multiple barcodes incorporated into their genome.

## Data and Software availability

NextClone and CloneDetective are available as open-source software on Github (https://github.com/phipsonlab/NextClone and https://github.com/phipsonlab/CloneDetective).

Documentation for NextClone and CloneDetective are available on https://phipsonlab.github.io/NextClone/ and https://phipsonlab.github.io/CloneDetective/respectively.

Code used to analyse the DNA-seq data and scRNA-seq data is available as a workflowr website on https://phipsonlab.github.io/NextClone-analysis/.

Raw FASTQ files for the DNA-seq are available on zenodo (https://zenodo.org/doi/10.5281/zenodo.10121942). BAM file for the scRNA-seq is available on zenodo (https://zenodo.org/doi/10.5281/zenodo.10129133 and https://zenodo.org/doi/10.5281/zenodo.10129624).

## Acknowledgements

This research was undertaken with the assistance of Milton HPC/Virtual Machines, supported by WEHI. We thank WEHI’s Research Computing Platform for providing advice and access to their High-Performance Computing facility.

