## Supplementary Material for "Extraction and quantification of lineage-tracing barcodes with NextClone and CloneDetective"

### Materials and Methods

For the 8k and 10k DNA-seq datasets, the MCF7 breast cancer cell line was transduced with the vexGFP ClonMapper library. Post transduction, cells exhibiting vexGFP expression were isolated, and subsequent cultures of 8,000 and 10,000 cells were established to form the respective 8k and 10k libraries. After an 18-day period of expansion, 1 million cells from each library were sampled and set aside for genomic DNA extraction. The barcode sequences were then amplified using PCR and the samples were then subjected to a targeted DNA-seq. The sequencing coverage for the 8k and 10k samples was 5.3X and 6.8X.

For the scRNA-seq data, the MCF7 cells were  barcoded following the ClonMapper protocol, processed using 10x Chromium protocol and subsequently sequenced. A total of 7,828 droplets were identified as cells, and the average library size per cell was 50,807, and the average number of genes detected per cell was 5,939.

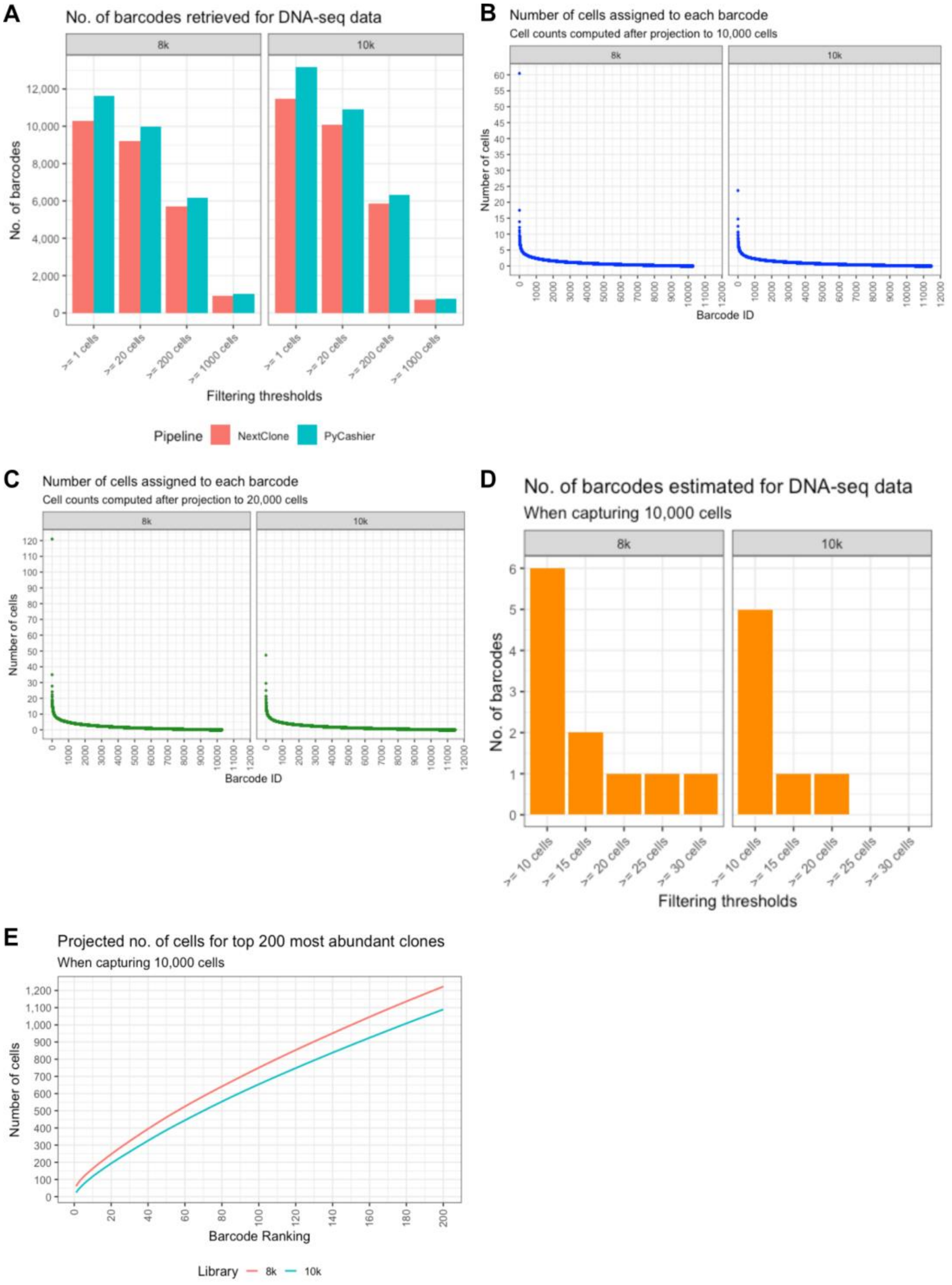


**Supplementary Figure S1.** Analysis of barcoded MCF7 cell line DNA-seq datasets with NextClone and CloneDetective. (A) Number of detected clone barcodes for 8k and 10k DNA-seq datasets obtained by NextClone and Pycashier at different filtering thresholds. (B) Projected cell counts per clone derived from the clone abundances for the 8k and 10k DNA-seq datasets when 10,000 cells are aimed to be captured for future scRNA-seq experiment. (C) Similar projection as (B) but for the scenario where 20,000 cells are captured. (D) Estimated number of clones which are represented by at least 10, 15, 20, 25 cells for 8k and 10k DNA-seq datasets when capturing 10,000 cells. (E) Cumulative number of cells representing the top 200 most abundant clones in the 8k and 10k DNA-seq datasets based on sequencing 10,000 cells.
